# A novel method for characterising the inter- and intra-lake variability of CH_4_ emissions: validation and application across a latitudinal transect in the Alpine region

**DOI:** 10.1101/2023.01.19.524063

**Authors:** Enrico Tomelleri, Katharina Scholz, Sylvie Pighini, Federico Carotenuto, Beniamino Gioli, Franco Miglietta, Ruben Sommaruga, Giustino Tonon, Alessandro Zaldei, Georg Wohlfahrt

**Affiliations:** Free University of Bozen/Bolzano, Faculty of Science and Technology, Bolzano/Bozen, ITALY; University of Innsbruck, Department of Ecology, Innsbruck, AUSTRIA; Institute of Bioeconomy - National Research Council - CNR-IBE, Florence, ITALY

## Abstract

Lakes in the Alpine region are recognised as critical CH4 emitters, but a robust characterisation of the magnitude and variability of CH4 fluxes is still needed. We developed a mobile platform for CH4 eddy covariance (EC) flux measurements to tackle this gap. Our approach was shown to be well suited to catch all CH4 emission pathways and overcome the limitations of other methods (e.g., gradient-based). This is by surpassing their local nature and thus being suited for characterising the variability of the within-lake emissions, primarily because of CH4 emissions by ebullition stochasticity. The mobile system was deployed at nine lakes across a latitudinal transect in the Alps and validated by comparing the measured fluxes with a fixed EC station and to chambers and boundary layer estimates. Methane fluxes were explained by water turbidity, dissolved organic carbon, dissolved nitrogen, elevation, particulate organic carbon, and total phosphorus. The highest fluxes and most substantial seasonal variability were found in a shallow low-altitude lake in the Southern Alps. Additionally, the mobile EC permitted to resolve the spatial structure of fluxes at the selected lakes. Finally, we demonstrated the usability of our novel mobile system to characterise intra- and inter-lake variability of fluxes. We suggest that characterising the intra-lake emission heterogeneity and a deeper understanding of inter-lake emission magnitude differences is fundamental for a solid estimate of freshwater CH4 budgets.

**Key Points:**

- CH4 emissions from alpine lakes are recognised to be an important component to the global methane budget but they are poorly characterized
- We developed and validated a mobile eddy covariance platform for capturing CH4 fluxes across lakes in the alpine region for two years
- A robust statistical model based on a few *in-situ* physicochemical and biological parameters can be generally used to predict CH4 fluxes

## 1 Introduction

Lakes are valuable sentinels of climate change (Adrian *et al*., 2009; Williamson *et al*., 2009; Moser *et al*., 2019) and are important in regulating the global water cycle. Temperature changes can significantly affect their phenology by modifying the seasonality of water temperature (Woolway *et al*., 2019). These changes can affect the trophic structure and the whole biogeochemical cycling; therefore, they feed back on the global climate system. In this context, lakes are increasingly recognised to contribute to the global methane (CH4) budget (Jansen *et al*., 2022; Matthews *et al*., 2020; Saunois *et al*., 2020).

Quantifying CH4 emissions is of key importance because they are recognised to contribute to offsetting the current terrestrial sink. The atmospheric CH4 concentration from a pre-industrial level of 800 ppb just broke the 1900 ppb mark (August 2022: 1908.61 ppb), with an average increase of 8.23 ppb/y over 2010-2020 and a maximum rate of 18.12 ±0.47 ppb in 2021 (Dlugokencky, NOAA/GML www.esrl.noaa.gov/gmd/ccgg/trends_ch4/). With a global warming potential over 100 years of 28 times CO2, CH4 plays a crucial role in the greenhouse effect (IPCC AR5). Second only to CO2, CH4 is estimated to have contributed 23% to the net anthropogenic radiative forcing during 1750-2005 (Etminan *et al*., 2016). Intensification of natural sources might result in up to 42% higher atmospheric concentration by 2100 (Christensen *et al*.,2019). According to Saunois *et al*. (2020), 47.5% of CH4 sources - in the period between 2000 and 2009 - are of anthropogenic origin, with significant contributions by agriculture and waste (57%) and the use of fossil fuels (33%). Natural sources account for the remaining 52.5%, with significant contributions from wetlands (40%) and freshwaters (43%). This last category includes lakes, ponds, reservoirs, and streams. The variability of their contribution is not completely understood at global scale, but it has been shown to have driven the anomalous growth in the most recent years (Peng *et al*., 2022).

While covering only a tiny fraction of the Earth s surface, freshwater ecosystems annually receive as much carbon (C) as the oceans (Cole *et al*., 2007), and much of this C can be buried and sequestered in the sediments (Downing *et al*., 2008). Because of their anoxic and highly reductive conditions, which promote methanogenesis, lakes are strong CH4 emitters. Other recent global estimates of CH4 emissions from freshwaters (Saunois *et al*., 2020) suggest a source of 159 Tg CH4 y-1 offsetting 25% of the total CH4 sink. This shows that freshwaters are crucial to the terrestrial greenhouse gas (GHG) balance. The same authors highlight that most open water emissions occur at a latitude between 25 and 54°N. Not only natural lakes but also hydropower reservoirs do release CH4, even though their emissions appear to have been overestimated in the past (Barros *et al*., 2011; Bastviken *et al*., 2011). A peculiarity of some hydropower reservoirs is the sudden release of CH4 accumulated in the hypolimnion due to the release of water through the turbine to the downstream rivers (Kemenes *et al*., 2007).

Estimating CH4 emissions of lakes and reservoirs is challenging because there are multiple pathways with diverse spatial and temporal dynamics. Such pathways are diffusion, ebullition, and plant-mediated transport (Bastviken *et al*., 2004). The CH4 production occurs in the anoxic sediment and the oxic water column (Bogard *et al*., 2014; Peeters and Hofmann, 2021). CH4 can then be transported to the surface either by ebullition or diffusion (Joyce, 2003; Liss & Slater, 1974), the first pathway mainly related to the anaerobic mineralisation of sediment organic matter (Aben *et al*., 2017; Segers & Kengen, 1998). The diffusive flux across the air-water interface is driven by the difference in water and atmospheric CH4 concentration and the piston velocity (Klaus, 2020). While ebullition results in a direct flux to the atmosphere, the portion of CH4 entering the water column and, potentially contributing to the diffusive flux, might be oxidised to CO2 by methanotroph bacteria which can eventually mitigate the CH4 flux from this pathway (Guggenheim *et al*., 2020). For stratified lakes, this pathway might be influenced by temporary CH4 storage in the anoxic layer, released by diffusion during lake overturn (Encinas Fernández *et al*., 2014). The fourth pathway includes plant aerenchyma mediated transport (Dacey and Klug, 1979; Henneberg *et al*., 2012) along the shores with emergent vegetation. This flux component is highly dependent on the vegetation characteristics and concurrent CH4 production/oxidation processes in the sediment.

Given the different factors affecting the emission pathways and their spatial and temporal heterogeneity, any estimate of the CH4 budget for a lake might be highly uncertain if not all pathways, location, and timing are accounted for. Currently, data-driven CH4 emissions synthesis studies are mostly biased toward the diffusion pathway (Engram *et al*., 2020). This is because the established underlying measurements are less capable of targeting ebullition. For example, the boundary layer method is based on the analysis of water samples. Therefore, the method is suitable for quantifying diffusive fluxes, but not ebullition.

Moreover, ebullition is a stochastic process because of its variability in time and space (DelSontro *et al*., 2011; Linkhorst *et al*., 2020) and bubble CH4 content (DelSontro *et al*., 2016). Therefore even estimates from lakes that have been intensively investigated utilising funnel flux traps are highly uncertain (Wik *et al*., 2013). The sampling designs associated with the different measurement methods are not necessarily constrained in time and space, making data-driven temporal and spatial up-scaling challenging. In addition, the method-inherent spatial and temporal representativity is highly dependent on the sampling strategy with constraints related to the number of samples per unit of space, time, and depth. The flux chamber diffusion method is also limited in spatial representativity because the chambers cover a limited surface and might randomly encounter ebullition. A network of automatic gas samplers (Duc *et al*., 2013; Bastviken *et al*.,2020) might be suitable for catching the temporal and spatial variability of diffusive fluxes, but one still might encounter difficulties in assessing the spatial patterns of ebullition. The eddy covariance (EC) method (Baldocchi, 2003) overcomes the other methods’ limitations by catching the spatial heterogeneity of all the CH4 emission pathways.

Additionally, it has the advantage of providing spatial integration across long time periods, being automated and nearly continuous. On the other hand, the measurement station typically anchored to a point in space (Baldocchi *et al*., 2020; Scholz *et al*., 2021), this method is very suitable for resolving the temporal dynamics within the so-called “footprint”, but it cannot be used for estimating the spatial heterogeneity across the whole area of interest. This issue can be addressed by installing the sampling system on a moving platform. The airborne EC method is a commonly used and widely accepted technique that has been validated against well-established tower-based EC (Gioli *et al*., 2004; Hiller *et al*., 2014; Vellinga *et al*., 2013). Applications for measuring CO2 fluxes from the ocean have been developed and validated (Miller *et al*., 2010; Butterworth *et al*.,2016; Dong *et al*., 2021). Still, shipborne EC measurements of CH4 fluxes remain occasional with only sparse disclosed methods and results (Thornton *et al*., 2020), which is probably related to the limited relevance of oceans in the global CH4 cycle compared with other ecosystems (Saunois *et al*., 2020). On the contrary, transporting EC equipment on lightweight boats offers a novel way to characterise spatial and temporal variability of CH4 fluxes from small and medium-sized lakes and thus for integrating the advantages of all the above-described methods.

In this study, we used an innovative mobile EC platform that allowed us to characterise the temporal and spatial variability of CH_4_ fluxes within and between lakes. The system consisted of EC instruments and sensors suitable for motion detection installed on a small boat. The whole system was lightweight and therefore transportable with a simple commercial boat trailer which allowed us to visit multiple lakes. While cruising on the lake, we also recorded micro-meteorological and bio-physical data. In addition, to validate our approach, we paired mobile EC measurements with a fixed EC tower, floating chambers, and dissolved gas samplings. Thanks to the portability, we made measurements in several natural and artificial lakes along a transect of two degrees of latitude from the southern to the northern side of the eastern Alps during the ice-free season in 2018 and 2019.

The objective of this study is thus to make use of our novel mobile EC platform to improve data availability and quality of CH4 emissions from lakes in regions susceptible to an increasing climatic variability like the Alpine region.

We tested the following hypotheses:

1. Higher temperatures are known to increase methanogenesis (Yvon-Durocher *et al*., 2011), while CH4 oxidation shows less dependency on temperature (Duc *et al*., 2010), and we thus expected CH4 emissions to decrease with elevation.
2. Higher substrate availability (dissolved and particulate organic matter) will result in higher CH4 emissions (Barros *et al*., 2011). We thus expected more productive lakes (more significant autochthonous C input) or lakes that have a larger net supply of allochthonous organic material to be characterised by higher CH4 emissions. This is expected to contribute to larger CH4 emissions from lakes located at lower elevation, which are usually fed by larger catchments, reach higher water temperatures, and are often more productive.
3. For the same size, relatively higher CH4 emissions are observed from shallow lakes because the distance from the sediment to the surface, along which CH4 may be oxidised, is shorter and because ebullition must overcome less hydrostatic pressure as compared to deeper lakes (Barros *et al*., 2011). Thus, we expected higher ebullitive emissions from shallower lakes, particularly if in combination with high substrate availability and temperatures, e.g., productive lakes at low elevations. Because hydropower reservoirs in the Alps are often built-in narrow valleys with steep slopes and are thus relatively deep, we expected, following (Diem *et al*., 2012), relatively lower CH4 emissions as compared to (more shallow) natural lakes at similar elevation.
4. Significant temporal variability in CH4 emissions exists in relation to the different phases of the thermal cycle such as ice-out (high emission due to accumulation of CH4 below ice cover), overturning (intense release due to convection of CH4-rich water followed by low diffusive emissions due to oxidative conditions thereafter), and lake stratification (high emissions due to oxygen depletion in productive lakes), as well as to variations in biological activity. Therefore, we expected to find an accentuated seasonality related to the lake physical and biological dynamics.
5. Compared to the widely employed floating chamber, funnel, and gradient methods, the eddy covariance technique provides spatially more representative measurements at a higher temporal resolution covering extended periods. We hypothesise then that this method will be capable of characterising the intra-lake temporal and the spatial heterogeneity of CH4 fluxes.

## 2 Materials and Methods

### Overview

We initially developed and implemented a CH4 eddy covariance flux equipment on a mobile platform: a 4 m aluminium boat (Marine Light 14M) operated with an electric engine (Minn Kota EO ½). We installed a u-shaped frame on the boat to support the micrometeorological equipment (Figure 1). This mobile platform was transported to nine selected lakes across the eastern Alps on a trailer and was used to conduct mobile CH4 flux measurements during the ice-free season. Roughly between May and October for the years 2018 and 2019. During each field campaign, water samples were taken and analysed for biological and physicochemical parameters. In addition, we characterised vertical profiles of physicochemical data *in-situ*. The EC fluxes were validated with a fixed EC station and compared with the boundary layer method and with chamber flux estimates.

**Figure 1:**
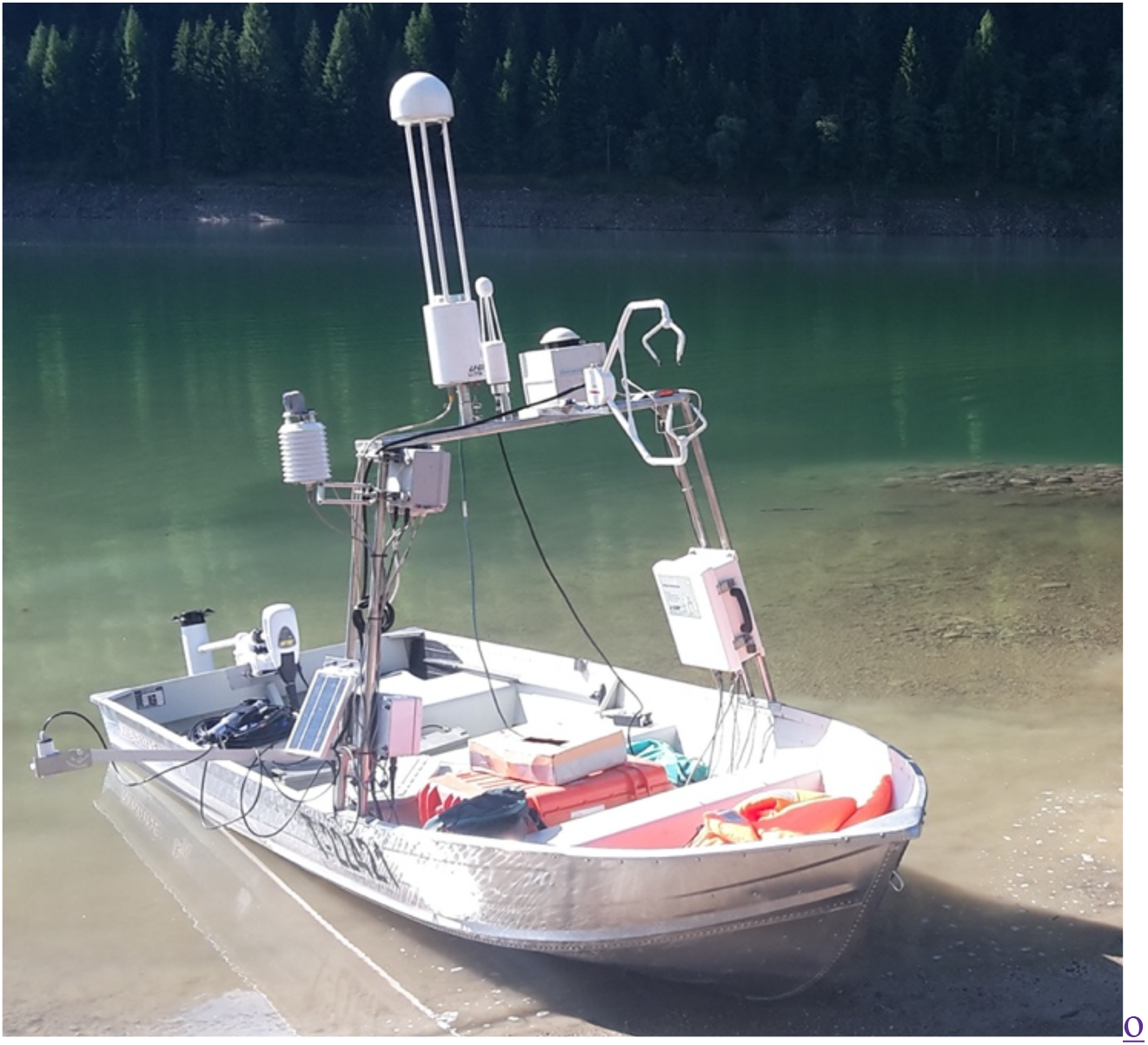
the boat equipped with instruments for eddy covariance measurements (a CSAT3 sonic anemometer and a LI-7700 open path CH4 analyser) and accurate movement detection (C-Migits Inertial Measurement Unit and a RT-20 ProPak Global Navigation Satellite System unit).

### Lake selection

Initially, a list of 76 lakes along a latitudinal gradient across the eastern Alps was compiled by extending a previous research by (Pighini *et al*., 2018). The lakes are located between 10.09° and 13.67° of longitude and between 45.79° and 47.88° of latitude (Figure S 1). Based on geophysical properties (elevation, area, shape, and depth) and lake origin (natural or artificial), the list was clustered through the k-means method (Hartigan and Wong, 1979) into 4 clusters (Figure 2). Differently from the other considered variables, “lake origin” is categorical, which required the use of the Gower distance (Gower, 1971) for computing the pairwise dissimilarities. This distance measure allows for computing the distance between entities with mixed categorical and numerical values. Cluster 1 includes lakes located at the outer latitudinal gradient. Cluster 2 includes shallow and small lakes in the southern part of the region of interest. Cluster 3 contains lakes at relatively high elevations, and cluster 4 mainly includes lakes of medium and large sizes located in the northern region at low elevations. From each cluster, we selected at least one lake to be visited during the exploratory phase resulting in the selection of a total of ten different lakes. Within-cluster selection occurred based on accessibility with the boat trailer. The Grünsee is not reachable with the trailer, but we still sampled it with other methods because of its altitude. Furthermore, we aggregated the fluxes of Heiterwanger See and Plannsee in our analysis because these two lakes connected by a 300 m long and 30 m wide channel.

**Figure 2:**
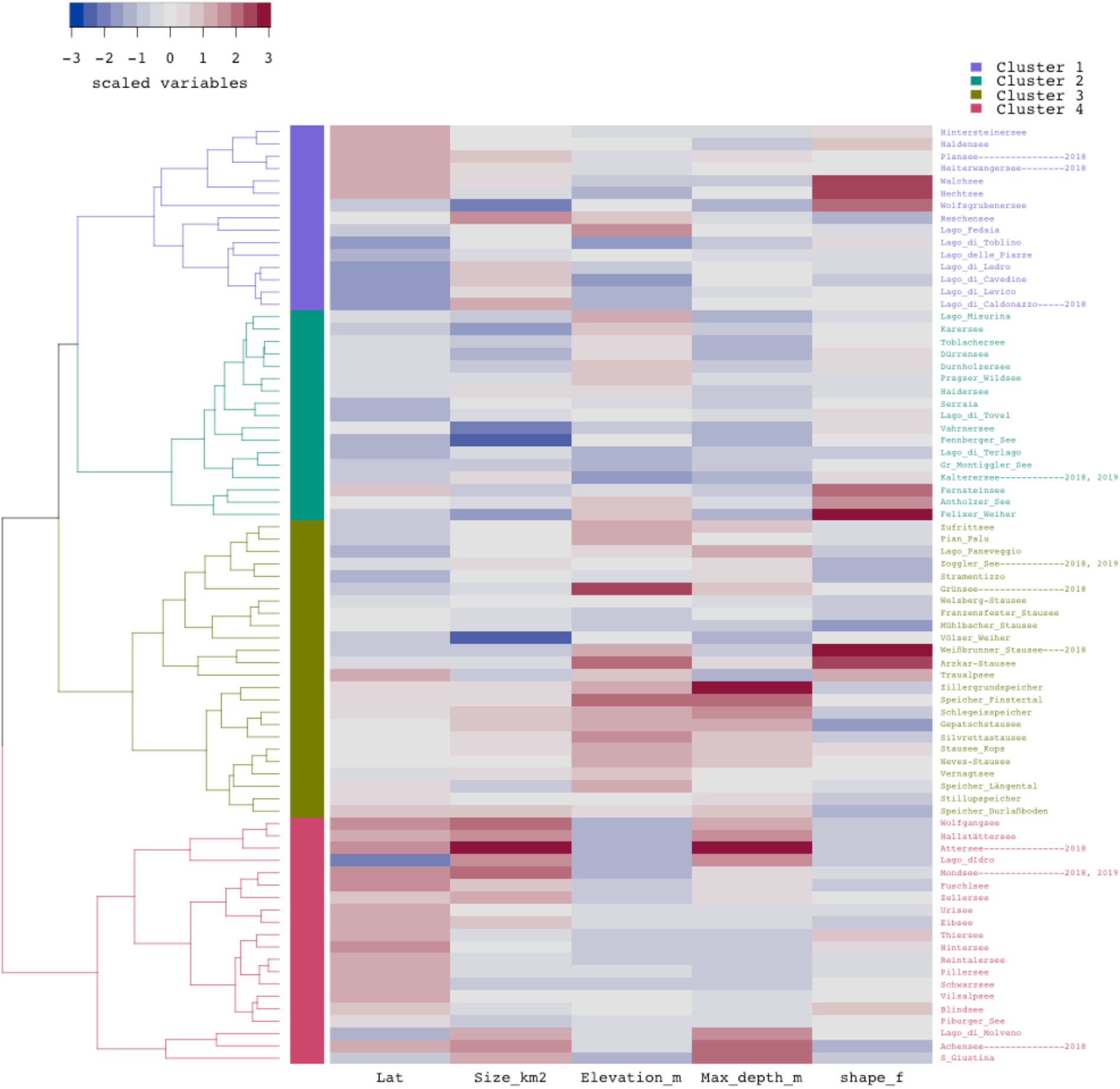
A hierarchical clustered heatmap showing lakes’ geographical and morphological characteristics in the region of interest. Sampled lakes and year of sampling are indicated on the right-hand side. Lat: latitude; Size_km2: surface area; Elevation_m: altitude above the sea level; Max_depth_m: lake depth; shape_f: ratio between the shorter and the longer axis.

The first field season in 2018 served to collect extensive information on the physicochemical properties and flux magnitude of the lakes. Based on this information, the pool of sites to be visited for intensifying the temporal sampling in the second field season was reduced to three in the second field season.

Firstly, the selected lakes were sampled between one and five times during 2018. These lakes ranged between 200-2500 m in elevation, 0.2-45 km^2^ surface area, and 6-170 m depth. Based on the results from the first study year, we selected three lakes (Mondsee, Kalterersee, and Zogglersee), which were sampled more intensively at regular intervals (4-6 times) during 2019. Mondsee was chosen because of the comparatively large surface area and depth and being intermediate in many of the physicochemical parameters (Tab. S1). Kalterersee was chosen because of the low depth and high concentrations of total P, dissolved N and C, chlorophyll-a and higher water temperature. Lastly, Zogglersee was chosen because it is a hydropower reservoir at high elevation and low temperature with low concentrations of total P, dissolved N and C, and chlorophyll-a.

Mondsee is an oligomesotrophic lake located in central Austria. The elevation is 481 m a.s.l. and it has a total area of 13.8 km^2^ with a mean depth of 36m and a maximum of 68m. The theoretical water exchange time is about 1.7 years (Dokulil & Skolaut, 1986). The main dischargers are the Griesler Ache and the the Zeller Ache. The northern sub-catchment has smooth hills generated by moraines based on Flysch rocks, while the southern sub-catchment is characterised by limestone and dolomite forming steep slopes (Kämpf *et al*., 2014).

Kalterersee is a eutrophic lake situated at a low elevation (216 m a.s.l.) in the Southern Alps. The total surface is 1.4 km^2^, and it reaches a maximum depth of 5.6 m (Balátová-Tuláčková *et al*., 1989). The watershed is relatively flat and small, and the outflow is low (mean outflow at the stream gauge “Grosser Kalterergraben” was 0.41 m^3^/s in 2019) which translates in a slow water exchange. The shores are covered by dense macrophyte vegetation.

Zogglersee is an oligotrophic artificial lake situated in the Rhaetian Alps at an elevation of 1141 m a.s.l. and with a total surface of 1.43 km^2^. The dam, which is 66 m high, was built between 1955 and 1964. The substrate is made of fluvioglacial soils, consisting of gravels and sand with boulders (Pagano *et al*., 2010). In 2018, the water level had a water level excursion (max-min) of 25.5 m and in 2019 of 30.8 m. The minimum level is typically reached in April, and the maximum is already in May because of snowmelt (Alperia s.r.l, p.c.). The water exchange time is highly variable as it depends on the snowmelt and the hydropower production.

### Description of sampling protocol

At each lake, the eddy covariance instruments were first installed on the boat. The inertial measurement unit was calibrated automatically by orientating the bow toward the magnetic north before powering the system. We made measurements considering most of the total lake surface but avoiding driving close to the shore. If possible, we steered the bow towards the main wind direction. Otherwise, we cruised crosswind, avoiding wind directly from the stern, which would cause biased measurements by the sonic anemometer due to flow distortion. We stopped at one random location during each cruise - away from the shore – to take three water samples for gas concentration and physicochemical and biological characterisation. Samples were kept refrigerated until the delivery to the laboratory for analysis within 1-2 days.

Additionally, we measured vertical profiles at approximately the same locations every 5 m depth with a multi-parameter probe (WQC24, DKK-Toa Corporation). These were repeated at two additional random locations. Only surface acquisitions were used for further analysis. During the second year of measurements, we also conducted chamber measurements for comparing our novel mobile eddy approach with this commonly used method (Deemer *et al*., 2016).

### Physicochemical and biological samples

Lake water was collected using three 1-liter plastic bottles at each sampling location. The bottles were filled entirely and closed with a screw cap about 10 cm below the water surface. In the laboratory, the samples were filtered through glass microfiber filters (Whatman; Glass Microfiber Filter; Quality GF/F). We performed different analyses for each filtered sample including total P, DOC, and the dissolved N, chlorophyll-a and C/N content by elemental analysis. We measured total P according to the Molybdate method (Vogler, 1966). We estimated DOC as non-purgeable organic C with a total carbon analyser (Shimadzu TOC-VCPH). At the same time, dissolved nitrogen (DN) was measured as total bound nitrogen (TN) with a Total Nitrogen Measuring Unit (Shimadzu TNM-1). For measuring particulate C (C_Filter) and N (N_Filter) content, we analysed the dried glass microfiber filters with the elemental analyser Flash EA 1112 (Thermo Electron Corporation) in the NC Soil configuration. Last, we estimated chlorophyll-a by analysing the filter residue of the water sample with a spectrophotometer according to (Lorenzen 1967) (Table S 1).

### Eddy covariance flux measurements

The boat (Figure 1) was equipped with a CSAT3 sonic anemometer (Campbell Scientific, Logan, UT, USA) and a LI-7700 open path CH4 analyser (LI-COR Inc., Lincoln, NE, USA) to measure wind velocity, wind direction, and atmospheric CH4 concentrations. The measurement height was about 1.4 m above the water surface. Eddy covariance flux measurements from floating and moving platforms need to account for the 3D movement of the wind sensor. Therefore, we additionally equipped our platform with a C-Migits Inertial Measurement Unit (IMU) (Systron Donner Inertial, Inc., Concord, CA, USA) capable of registering pitch, roll, and yaw at a frequency of 10 Hz in a fashion described in (Vellinga *et al*., 2013). The instantaneous speed of the sonic anemometer (i.e., speed and direction of the boat) was obtained employing an on-board RT-20 ProPak Global Navigation Satellite System unit (GNSS) (NovAtel Inc., Calgary, Canada). We synchronised timestamps using a unique standard time reference system during the data logging procedure. The measured wind signal was then rotated and corrected according to the IMU and GNSS data. The measurement units for recording motion were installed on the same horizontal plane as the micrometeorological instruments; therefore, the vertical displacement was negligible for the motion corrections.

CH4 fluxes were calculated based on the motion-corrected wind vectors and the recorded CH4 molar density using the software EddyPro 7.0.6 (LI-COR Inc., Lincoln, NE, USA) following established procedures (Aubinet *et al*., 2000; Aubinet *et al*., 2012). Raw data processing included de-spiking, double rotation of the wind data, WPL corrections, and the detection and compensation of time delays in the CH4 data relative to the wind signal. CH4 fluxes where then derived from the covariance of the vertical wind speed and the CH4 concentration. We averaged the covariances over 5-minute intervals to obtain a high temporal (and as the boat is moving also high spatial) resolution of flux estimates. Commonly, fluxes are calculated for 30-minute intervals. In our case, shorter averaging times are justified by the low measurement height. We corrected spectral losses in the low and the high frequency range according to (Moncrieff *et al*., 1997; Moncrieff *et al*.,2004). We labelled the quality of aggregate fluxes with a scale from 0 to 2 according to (Foken *et al*., 2004) where “0” is the best quality, “1” indicates that data are suitable for general usage while “2” indicates low quality data.

Additionally, we identified the source area of the measured signal by applying the footprint model proposed by Kljun *et al*. (2015). The footprint model was superimposed on a land-water mask to determine the underlying surface using the web mapping service Bing Maps. For further analysis, we only kept fluxes when the respective footprint was more than 90% on the water surface.

### Boundary layer method

During the campaigns in 2018 and 2019, we collected water samples for dissolved gas analysis congruent with our EC cruises. By applying the gradient method, we could estimate CH4 fluxes. We collected water probes by sampling surface water with a manual sampler (Guérin *et al*., 2007). In doing this, we filled 60 mL brown glass vials with the water samples by letting it flow in until about twice the volume of the vial. To prevent any biological activity from affecting the sample, we added copper chloride (as described in Diem *et al*., 2012) before sealing the vials with a rubber cap and a metal crimp. The collected samples were then preserved upside down in the dark until laboratory analysis. In the laboratory, preceding the analysis, we created a headspace in each vial by displacing about one-third of the total sample volume with nitrogen as described in (Guérin *et al*., 2007). Then each vial was vigorously shaken to equilibrate dissolved CH4 between water and gas phases. We performed the gas analyses with an Agilent 7890A Gas Chromatographer (GC) provided with a Flame Ionization Detector (FID) for CH4. For each water sample, we injected twice 0.5 mL of the headspace gas into the GC using a gastight syringe. If the difference between the two injections was higher than 5%, then a third injection was made. We used a commercial gas standard mixture (CH4 at 8.11 ppm, CO2 at 501 ppm and N2O at 903 ppb) for calibrating the GC before analysing the samples, and every 10 samples. The dissolved gas concentrations were estimated then according to Weiss (1974).

To estimate the air-water gas transfer we made use of the gradient method. The method is based on the idea that the primary driving mechanism that regulates the gas transfer velocity (k) across the air-water interface is the near-surface turbulence. Under the assumption that wind speed largely drives near-surface turbulence at lakes, we estimated k based on measured wind speed. To that end, we firstly scaled wind speed from the sonic anemometer to wind speed at 10 meters height (U10) using the logarithmic wind law. The value of the gas transfer velocity for CH4 normalised to a Schmidt number of 600 (k600; i.e., at a standard water temperature of 20 °C) was obtained using the model described by (Crusius & Wanninkhof, 2003) which accounts for a bilinear dependence between k600 and U10. Secondly, we converted k600 to k by accounting for the measured water temperature during the sampling. Thirdly, we obtained the CH4 flux by multiplying k with the difference in gas concentration between water (from GC) and air (from the CH4 analyser on the boat) as indicated by (Raymond *et al*., 2012).

### Floating chambers

The chambers consisted of a plastic container with a 0.36 m radius and 0.14 m height. The container was supported by a floating body and shielded with reflective material to limit solar radiation effects on internal temperature. Each chamber was fitted with a rubber septum on its top for gas extraction and a valve to equilibrate pressure during the initial positioning. After the chamber’s placement on the water surface the valve was closed, and we extracted an initial gas sample through the rubber septum. Subsequently, we extracted 3 additional gas samples at 10-minute intervals for a total of 30 minutes measurement period. Each gas sample was injected in an evacuated 2 ml vial capped with a massive butyl rubber stopper. We used two well distanced floating chambers at each sampling location for parallel measurements. Finally, CH4 concentrations were obtained with the same analytical process as for the dissolved gas samples. The rate of change across the four samples of each measurement series was utilised to calculate the flux between water and air as described by (David Bastviken *et al*., 2010; Podgrajsek *et al*.,2014).

### Fixed tower validation

To validate the CH4 fluxes measured by the mobile system on the boat, we compared those flux estimates to fluxes obtained by a fixed EC tower situated on the north-western shore of Mondsee. The tower system was operative from April 2019 to October 2020. Firstly, we estimated tower CH4 fluxes for matching time intervals (i.e., 5-minutes intervals during times when also measurements with the boat EC were conducted on Mondsee). Secondly, we identified the boat’s exact location during those times and estimated position and shape of the instantaneous footprints according to (Kljun *et al*., 2015). Likewise, footprints were calculated for the fixed EC tower measurements and the overlap between the tower and boat footprints was analysed (Figure 3).

**Figure 3:**
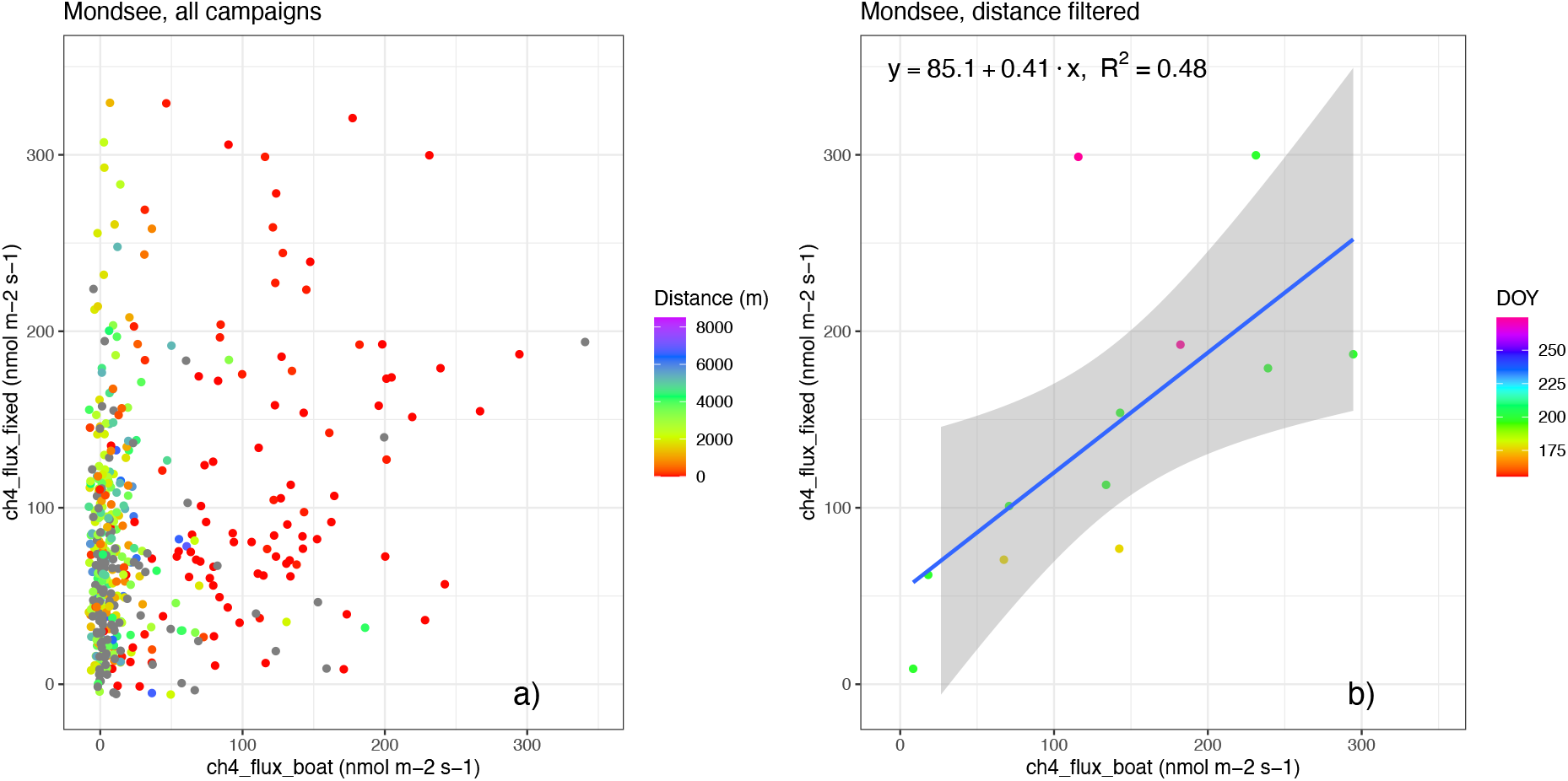
Comparison of a) all CH4 fluxes measured from the shore and the boat by EC. The color scale indicates the distance of the boat from the fixed station; b) CH4 fluxes for matching footprints (overlap > 75%). The color scale indicates the day of the year of the respective measurement campaign.

### Statistical analyses

All statistical analyses were conducted with the statistical software R (version 4.0.2). Firstly, a validation of measurements with the mobile system is presented. We used a simple linear regression to compare our results with the fixed tower located at Mondsee. The predictive power was evaluated by including further explanatory variables in a multiple regression. Linear regression slope was tested with the Pearson correlation coefficient. The principal component analysis (PCA) was performed by a singular value decomposition of the centred and scaled data matrix. To understand the spatial heterogeneity of CH4 fluxes measured with the EC method, we deployed the Moran I metric (Bivand & Wong, 2018; Moran, 1950) on the 5-minute aggregated fluxes. Moran’s I is a measure of spatial autocorrelation. Values of the Moran I metric not significantly different from 0 show perfect randomness, negative values indicate a spatial dispersion, while positive values mean clustering of spatial patterns.

## 3 Results and discussion

### Method Validation

We found a weak correlation between the CH4 flux measurements of the mobile EC and the fixed system at Mondsee when all available flux pairs were included (Figure 3a). With a closer inspection, by i) filtering CH4 fluxes from multiple campaigns for data quality (QC) lower than 2, ii) selecting footprint overlap higher than 75%, and iii) separating the distance between boat and tower lower than 100 m, we could confirm that the CH4 fluxes from the mobile station corresponded to the fluxes measured with the fixed EC system on the shore (R2 = 0.48, p-value: 0.00577) (Figure 3b). On the other hand, a multiple regression on the filtered dataset, including the horizontal distance between the systems and the footprint overlap, showed higher explanatory power (R2 = 0.7, p-value: 0.0056). This confirmed that lower distance and higher overlap favor the CH4 flux comparison among the different measuring systems. These results demonstrated the validity of our mobile system for conducting CH4 measurements.

To further validate the CH4 flux estimates from our mobile EC system, we compared our results with more established approaches such as the boundary layer method and floating chambers. The correlation between the results of the boundary layer method and the mobile EC system are consistent (slope 0.81, Figure S 2). The magnitude of CH4 fluxes estimated with the mobile EC across lakes was not significantly different from that estimated by the boundary layer method (linear regression slope not significantly different from 1 with p = 0.178). Similar results were also found for the chamber approach (slope 0.87), conducted during the second year of field activities (Figure S 3). While other authors (Erkkilä *et al*., 2018) found the gradient-based method to give mainly lower flux estimates of CH4 fluxes compared to the EC method, we cannot find significant differences at whole-lake scale across all the sites of our study. This confirms that the average fluxes are not method dependent, partly in contrast with findings by (Sanches *et al*., 2019). However, in the specific case of Kalterersee, we found higher fluxes with the EC method than the boundary layer method (Figure S 2) and the chamber method (Figure S 3). We hypothesise that, contrary to the other methods, the EC approach can systematically capture CH4 emissions from ebullition. Therefore, significant differences among flux estimates using different methods can be expected in lakes where this pathway is relevant, and EC remains the only method capable to include all pathways.

### Variability of CH4 fluxes

The largest CH4 emissions, with a pronounced seasonal course, were consistently measured at Kalterersee (Figure 4), exhibiting the largest within-campaign variability. It showed an average flux of 154.6 nmol m^-2^ s^-1^. The measured CH4 fluxes at Kalterersee were consistent with those values measured with the EC method by (Iwata *et al*., 2018) at Lake Suwa (JP) - up to 151 nmol m^-2^ s^-1^ - which is a shallow and eutrophic lake and with those measured by (Waldo *et al*., 2021) at Lake Acton (US) - up to 302 nmol m^-2^ s^-1^ - which is a small hypereutrophic reservoir. Such high CH4 fluxes are likely related to shallowness, the high temperature in summer combined with the high organic matter flow from the vegetated shores. The flux seasonality is reflected in the temporal variability of surface water temperature during the measurement period, which exhibited an interquartile difference of 8.9 °C ranging from 15 °C (14 May 2019) to 27.9 °C (8 July 2019). This corroborates our results from a correlation analysis of the whole dataset in which we found that CH4 fluxes to correlate positively with water temperature and negatively with maximum depth (Figure 5).

**Figure 4:**
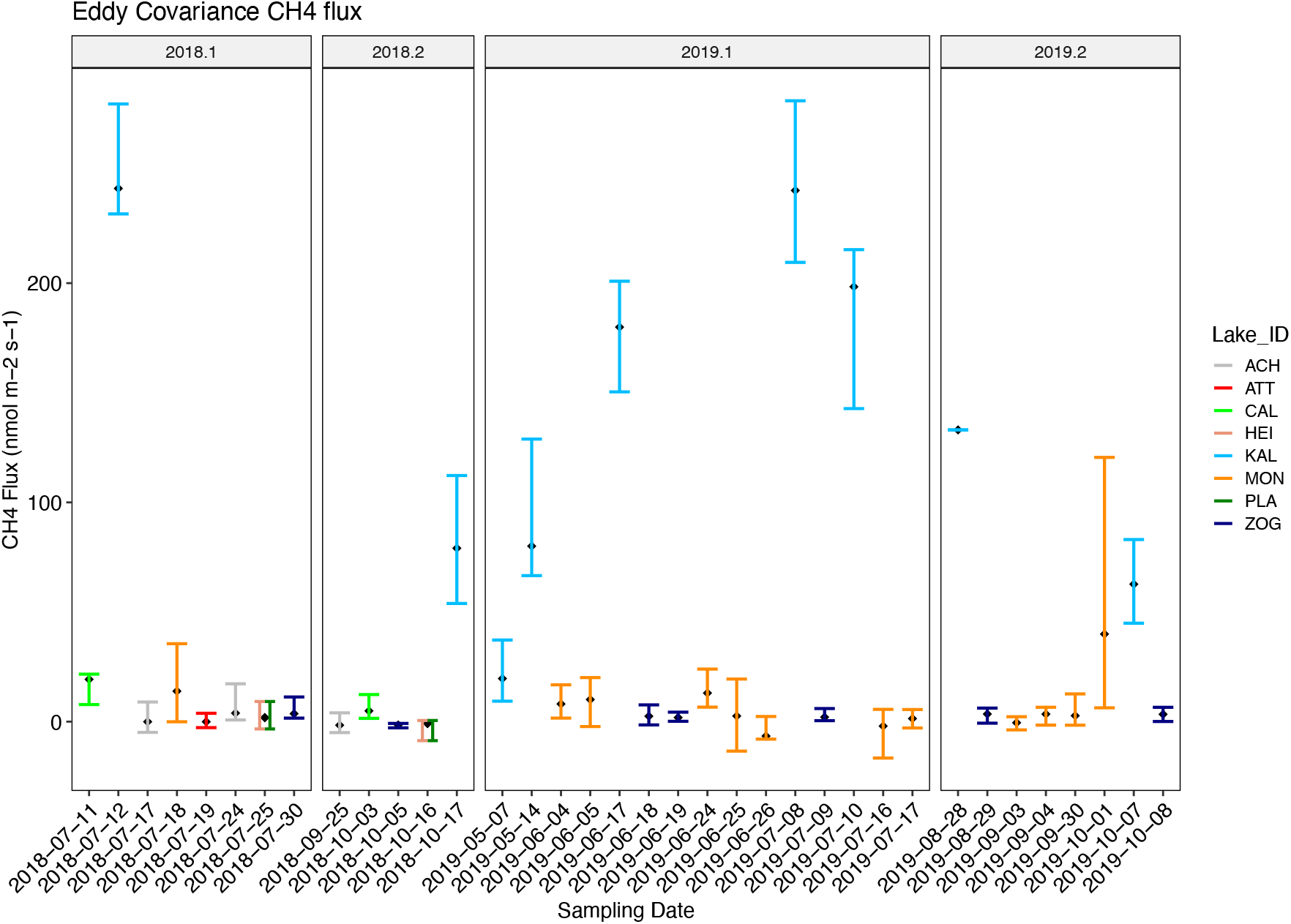
Temporal dynamics and spatial variability (interquartile difference) of EC CH4 fluxes for each field campaign during the measurement period (2018 and 2019). The boxes labelled with x. 1 - where x is the measurement year - depict the spring and summer campaigns, while the boxes labeled with x.2 depict the autumn campaigns. ACH: Achensee; ATT: Attersee; CAL: Lago di Caldonazzo; HEI: Heiterwangersee; KAL: Kalterersee; MON: Mondsee; PLA: Plansee; ZOG: Zogglersee. The frames 2018.1 and 2019.1 represent the summer campaigns while the frames 2018.2 and 2019.2 the autumn campaigns.

**Figure 5:**
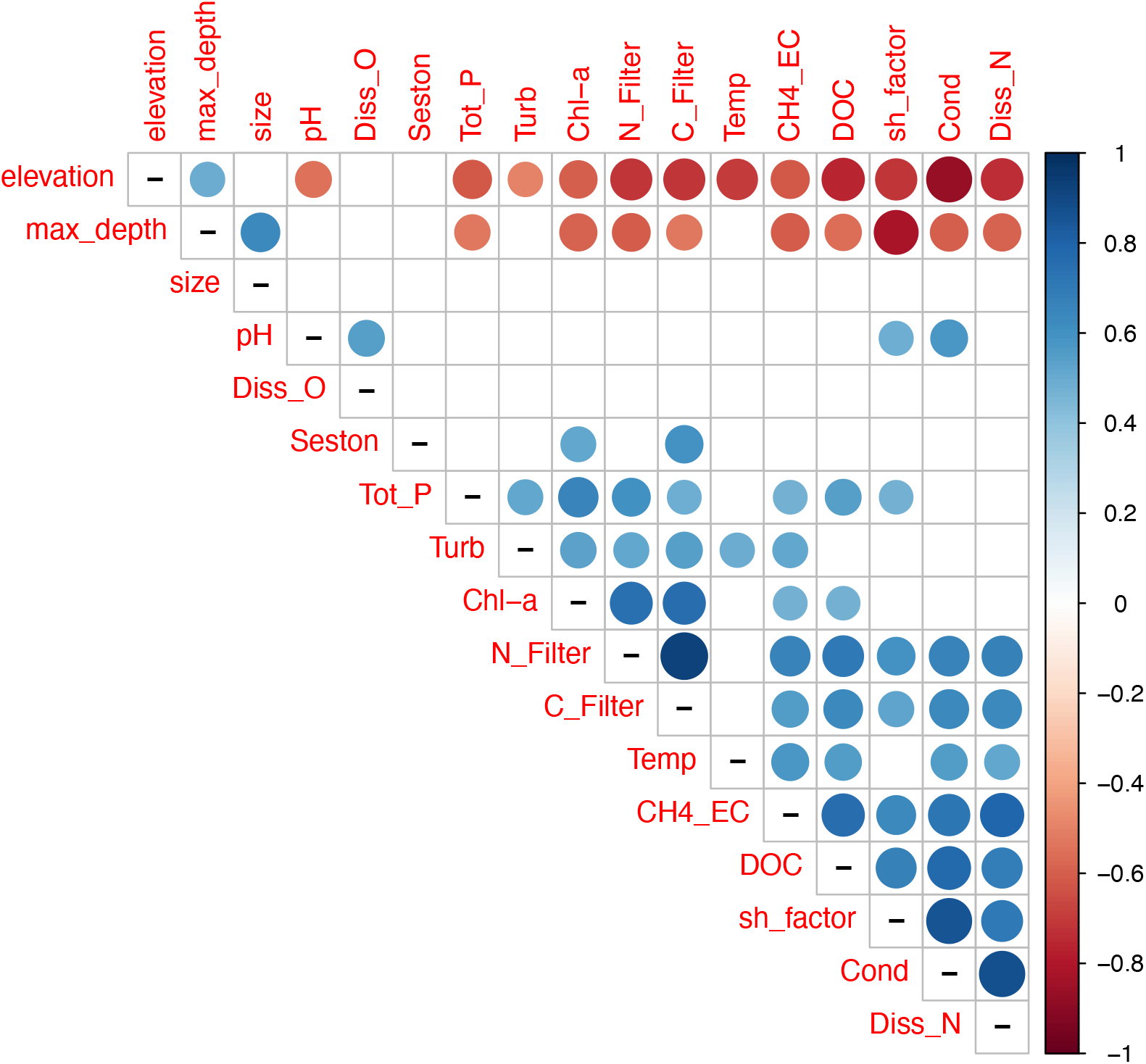
Correlation analysis of physical, biological, morphological characteristics, and mean eddy covariance CH4 flux for each field campaign. Only significant correlations (p < 0.01) are shown. The correlation matrix is ordered according to a hierarchical clustering of the correlation coefficients. Abbreviations: elevation, altitude above the sea leve (m); size, lake surface (ha); Max_depth, lake depth (m); pH, power of hydrogen; Diss_O, dissolved oxygen (mg/l); Seston, C:N ratio of suspended particulate matte; Tot_P, total phosphorus (μg/l); Turb, turbidity (NTU); Chl-a, chloropyll-a content (μg/l); N_Filter, particulate nitrogen (μg/l); C_Filter, particulate carbon (μg/l);); Temp, temperature (°C); CH4_EC, CH4 flux measured with the EC method (nmol m-2 s-1); DOC, dissolved organic carbon (μg/l); sh_factor, ratio between the shorter and the longer axis; Cond, conductivity (mS m-1); Diss_N, dissolved nitrogen (μg/l); Cond, conductivity; Tur, turbidity (NTU; Sal, salinity), DOC, dissolved organic carbon (μg/l); DN, dissolved nitrogen (μg/l); Seston, C:N ratio of suspended particulate matter.:.

The second largest emitter was Mondsee with an average flux of 7.5 nmol m^-2^ s^-1^ and with the peak emission found in June/July. The measured fluxes showed a lower spatial variability than Kalterersee but were still larger than the other lakes. The spatial variability is normally assumed to be related with lake size, the heterogenous land cover at the shores, and the depth variability (Schilder *et al*., 2013). At this lake, the shore of the north-western arm is more urbanised, whereas the south-eastern arm is significantly deeper. Additionally, the Griesler Ache, the primary tributary and source of suspended sediments, flows into the lake from the west at a perpendicular angle to the outflow direction (Kämpf *et al*., 2014). This is possibly accelerating the outflow in the southern half of the lake and therefore causing a gradient in sediment concentration and water temperature.

Zogglersee showed the lowest magnitude and variability of CH4 fluxes with an average value of 2.3 nmol m^-2^ s^-1^. Unlike Mondsee, where floods occur mainly in summer - during the productive period - the peak inflow at Zogglersee occurs during snowmelt (i.e., the peak lake level is typically reached in May). Therefore, transport of organic sediment into the lake is probably lower than in Mondsee. Further, the lake low water temperature probably limits methanogenesis. Seasonal changes of water temperature at Zogglersee were moderate with an interquartile difference equal to 3.65 °C (maximum temperature 18.1°C on 29 August 2019). The fluxes from these last two lakes were consistent with those reported for lakes of similar characteristics such as L. Kuivajärvi (5.9 nmol m^-2^ s^-1^, Erkkilä *et al*., 2018), but were lower than the average value of 28.21 nmol m^-2^ s^-1^ reported by (Johnson *et al*., 2021) for boreal reservoirs. This difference might be related to the higher organic matter content of such lakes and partly related to the high elevation/lower temperature of Zogglersee.

In all lakes, we occasionally measured negative fluxes, meaning that the flux of CH4 from the air to the water was dominant. These fluxes could be generated by diverse concurrent random circumstances (Prytherch *et al*., 2015).

### Drivers of CH4 fluxes

We compared the measured biological parameters with those measured for 207 boreal lakes by (Juutinen *et al*., 2009). Generally, we found that the lakes in our study generally have a higher pH (8.13 vs 6.6) and higher turbidity (4.41 vs 1.5 NTU), but lower phosphorus concentration (4.67vs 16.0 μg/l). Total organic carbon was lower (2.8 vs 9.0 mg/l), while total nitrogen was comparable (557 vs 505 μg/l). The measured CH4 concentrations in the surface water (0-6 μmol/l) were comparable with values reported for Finnish lakes, 0.2-1.8 μmol/l (Juutinen *et al*., 2009), with oligotrophic Swedish lakes, 0.1-1.9 μmol/l (Bastviken *et al*., 2004), with Wisconsin lakes, 0.3-2.3 μmol/l (Bastviken *et al*., 2004) and with a previous study in the same region, 0-5.9 μmol/l (Pighini et al., 2018).

By scrutinising the results from the biological analysis and the geophysical properties by using linear correlations, we found that CH4 fluxes measured with the mobile system negatively correlate with elevation and maximum depth (Figure 5). On the contrary, CH4 fluxes were positively correlated with water temperature, total phosphorous, turbidity, chlorophyll-a, C and N content. Such parameters are related to sediment fluxes and biological activity, which are demonstrated to link with methanogenesis (West *et al*.,2015; Yakimovich *et al*., 2020). The correlation of CH4 fluxes with temperature was consistent with the findings of (Duc *et al*., 2010), supporting the hypothesis that methanogenesis and oxidation are not equally responding to temperature changes.

A stepwise model selection by the Akaike information criterion (AIC) indicated that turbidity (Tur, p < 0.001), dissolved organic carbon (DOC, p < 0.001), dissolved nitrogen (DN, p < 0.001), elevation (Elevation_m, p < 0.01), dried filter carbon (C_filter, p < 0.1) and total phosphorus (TP, p < 1) can predict average lake CH4 fluxes as measured from our EC system (R2 = 0.87). This is consistent with the findings of CH4 concentration of five montane lakes in the Sierra Nevada by (Perez-Coronel *et al*., 2020) and, in particular, the dependency of the CH4 fluxes magnitude on the elevation of the lake/catchment as initially hypothesised.

Contrary to (Bastviken *et al*., 2004), both with single and multiple regressions analysis, we could not find any clear correlation between CH4 emissions and lake size. This might be related to the fact that the size effect is marginal compared with other factors included in our study. In addition, we did not find any significant correlation with pH. This indicates that pH values observed within our lakes (provide observed range) do not limit methanogenesis, as reported by (Bastviken 2009). Additionally, dissolved oxygen could not explain any variation in the CH4 fluxes. We assume that this is because the surface water is usually well oxygenated during the ice-free season, and this measurement does not provide information about methanogenesis in the deeper and possibly anoxic layers or in the sediments. Lastly, (Bastviken *et al*., 2003) show that methanotrophic bacteria can be a substantial food supply for zooplankton. Therefore, we expected that seston - which comprises all biological components - could have had an explanatory power for CH4 fluxes, but we could not confirm this.

To further understand the drivers of CH4, we performed a PCA. We used average CH4 fluxes for each campaign and the *in-situ* correspondent parameters (Figure 6a). We then replicated the PCA using the parameters obtained with laboratory analysis (Figure 6b). In the first case, we found that CH4 fluxes contributed to the spread along the principal component (45.4% explained variance) together with conductivity (Cond) and temperature (Temp). Dissolved Oxygen (Diss_O), pH, temperature, and turbidity contributed to the spread along the second component (30.6% explained variance). While the spread along the first component reflected the inter-lake differences, the intra-lake differences were to be resolved in the second component. This can be explained with the moderate (Mondsee) and strong (Kaltersee) CH4 emission seasonality of lakes that are situated at the extremes of our latitudinal gradient. In fact, it was also evident that the autumn campaigns were situated at the chart’s top while the spring ones were at the bottom. We assumed that this was caused mainly by the temperature seasonality.

**Figure 6:**
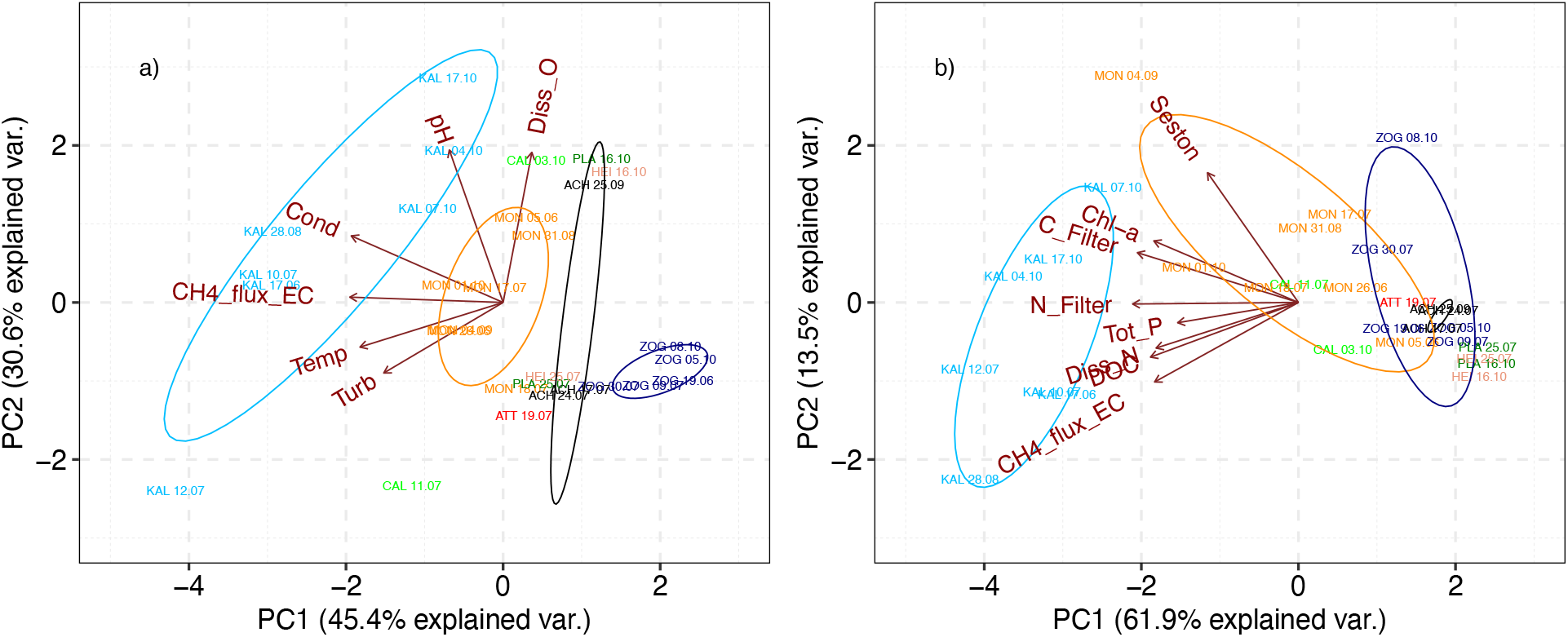
Principal Components Analysis of a) in-situ parameters and average EC CH4 fluxes and b) laboratory analyses and average EC CH4 fluxes. Samples are displayed as symbols while variables are displayed as vectors.

By analysing the principal components of CH4 EC fluxes and laboratory analysis (Figure 6b), we confirm the contribution of all the parameters along the first component (61.9% explained variance), which reflects most of the differences between lakes, but little of the within lake temporal variability. This might be related to the fact that the magnitude of such parameters within a lake is strongly affected by the intrinsic biophysical properties of a specific site. Their heterogeneity among lakes might mask the within-lake temporal variations.

### Spatial structure of CH4 fluxes

Our mobile EC system permitted us to measure the CH4 fluxes heterogeneity at each site by geolocating the 5-minutes aggregated flux estimates (Figure 7a, 7c, 7e). We could not identify any spatial pattern in the CH4 fluxes from a visual interpretation of the maps. By using the Moran I metric as a measure of spatial autocorrelation for the 5-minutes aggregated fluxes, we could identify some differences among the three more intensively investigated lakes. At Zogglersee, we found that the Moran I metric is not significantly different from 0 at any distance (Figure 7f). This means that the fluxes do not have a spatial structure, and since they are uniformly low, we assumed that the main emission pathway was diffusion. This contrasts with the findings of (Beaulieu *et al*.,2020) for diffusive and ebullitive emission rates from 32 reservoirs in North America. In fact, the authors report that ebullition being the dominant emission pathway in 75% of the systems. On the other side, (Sollberger *et al*., 2017), by deploying a floating eddy covariance system over two years at a reservoir in the Alpine region, confirmed that the fluxes were uniformly low. The presence of ebullition would cause some heterogeneity and thus we can assume that the primary pathway was diffusion. The physical characteristics of Zogglersee are very similar to the studied lake by (Sollberger *et al*., 2017). Therefore, we can hypothesise that most of the high altitude and deep reservoirs are CH4 emitters with low heterogeneity. At Mondsee, we found positive Moran I metric values between 0 and 400m of distance (Figure 7d) from each geolocation (5-minutes aggregated fluxes). This indicates that the spatial distribution of high values and/or low values in the dataset is more spatially clustered than expected if underlying spatial processes were random. This indicates the presence of gradients which we assume are influenced mainly by the bathymetry. In fact, Mondsee is shallower at the northern shore than the rest of the lake (Kämpf *et al*., 2014). At longer distances, the spatial structure becomes random until 4.8 km. In the range between 4.8 and 6.4 km, we found negative values significantly different from 0, which means the spatial distribution of high and low fluxes at this scale is more spatially dispersed than would be expected if underlying spatial processes were random. Such distances reflect the differences in the northwestern and south-eastern sections of the lake. These are mainly related to the effects on water exchange and sediment caused by the main tributary, which flows into the lake from the middle of the west shore (Kämpf *et al*., 2014). This shows that site-specific hydrological dynamics are to be accounted for when evaluating the CH4 budget of a lake. Lastly, the Moran I metric for Kalterersee was found to be negative up to 100 m of distance (Figure 7b). This shows that the fluxes were perfectly dispersed as it would be expected for dominance of the ebullition CH4 emission pathway. This can be explained by the combination of two factors: by the uniform shallowness combined with the abundant aquatic and underwater vegetation. These are both factors which have been already identified as control on magnitude and spatial distribution of ebullition fluxes (Wik *et al*., 2018).

**Figure 7:**
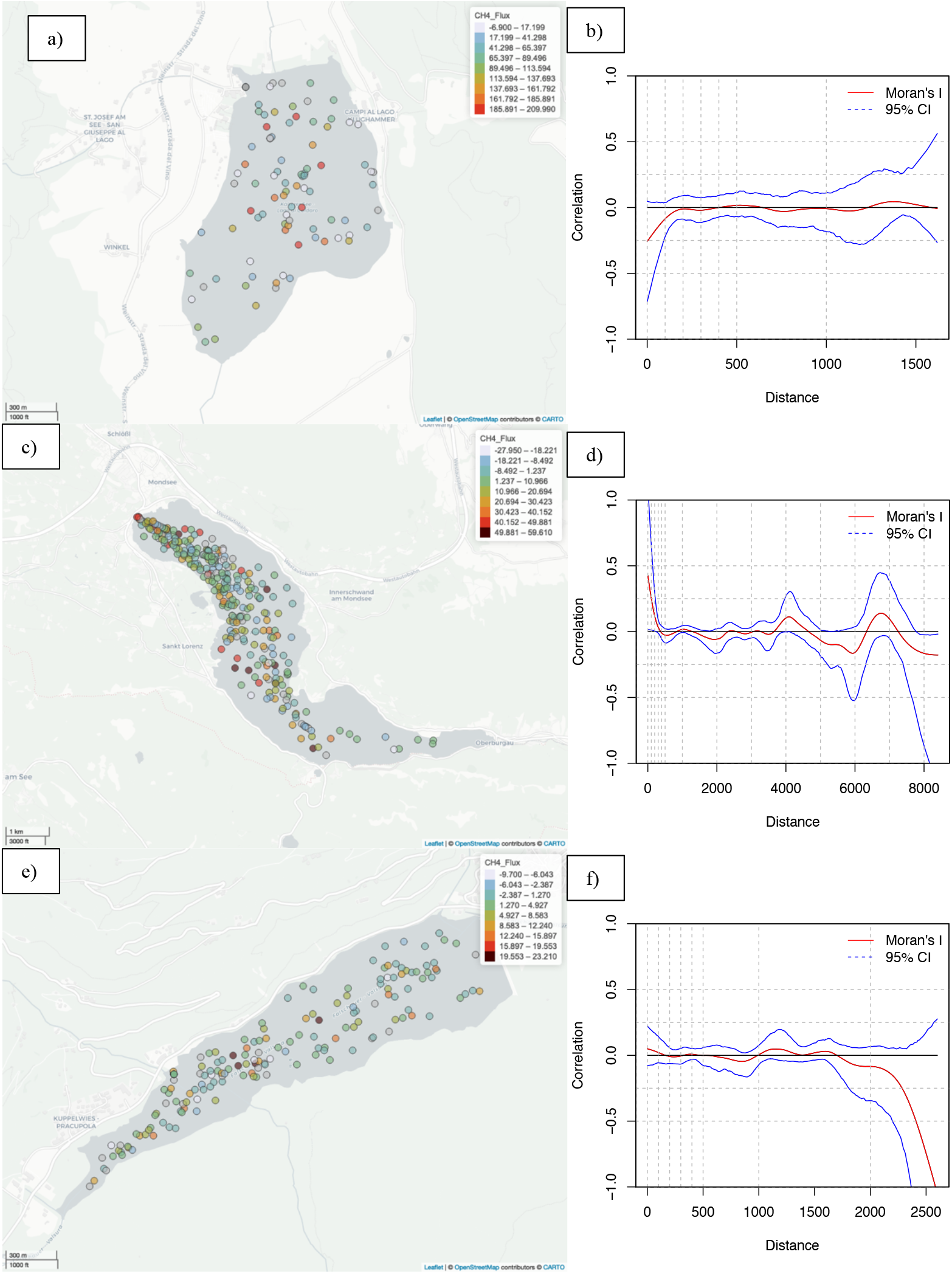
Maps of CH4 fluxes across all the measurement cruises for a) Kalterer See, c) Mondsee and d) Zogglersee and b), d), f) relative spline-correlograms. The confidence interval of the Moran I metric was generated with bootstrapping.

## 5 Conclusions

A mobile EC system was demonstrated to be suitable for characterising the CH4 emissions from a lake including the characterisation of their temporal and spatial heterogeneity. The system was utilised at 9 different lakes across the eastern Alps. The ancillary data collected during the cruises were used to explain the drivers of CH4 emission. Firstly, we could demonstrate that CH4 emissions decrease with elevation. In addition to this we found higher emissions from the shallower lake, particularly if in combination with higher temperatures, e.g., lakes at lower elevations. Thus, we could confirm our first hypothesis. At the same time, we could also confirm our second hypothesis that higher substrate availability results in higher CH4 emissions. In fact, we could find significant positive correlations with total phosphorous, turbidity, chlorophill-a, particulate nitrogen, and carbon thus more productive lakes (more significant autochthonous carbon input) or lakes that have a larger net supply of allochthonous organic material were characterised by higher CH4 emissions. This translates into more emissions for warmer autotrophic lakes. In addition to this, we found that for the same size, relatively higher CH4 emissions are observed from shallow lakes because the distance from the sediment to the surface, along which CH4 may be oxidised, is shorter and because ebullition must overcome less hydrostatic pressure as compared to deeper lakes. This confirms our third hypothesis. Fourthly, we could not confirm an accentuated seasonality related to the lake physical and biological dynamics for all the studied lakes. On the other hand, the warmer, shallower, and heterotrophic lake Kalteresee confirmed our expectations showing the maximum of emissions for the month of July. This was seen for both the measurement years. Because of the geomorphological characteristics of this lake, we can assume that such a behaviour is not related to stratification but rather to variations in biological activity. Anyhow, we believe that extending the measurements could help to better characterise the seasonal CH4 dynamics for other lakes and detect hot moments in time like the ice-out or the overturning. Lastly, we demonstrated that - compared to the widely employed floating chamber, funnel, and gradient methods – the eddy covariance technique from a mobile platform provides spatially more representative measurements at a higher temporal resolution covering extended periods. Thus, the approach showed to be suitable for characterising the intra-lake and inter-lake variability of CH4 fluxes. By acquiring ancillary biophysical information, we could characterise which parameters are suitable for describing the between lakes and between days variability. In fact, a robust statistical model based on few *in-situ* environmental parameters can be generally used to predict CH4 fluxes. Such characterisation can be deployed for upscaling our results and be used for classifying the emission of lakes taking advantage of the results from a limited number of time demanding and data rich field campaigns. This step is crucial for an accurate quantification of sources especially in the context of a warming climate where the effect on this specific ecosystem type could have relevant positive feedbacks on the destabilisation of the earth’s climate system.

## Acknowledgements

This research was funded by the Autonomous Province of Bozen/Bolzano (ALCH4 project). Gry Larsen is thanked for conducting the laboratory water analyses; prof. dr. Andreas Richter is thanked for providing the Li-7700 for CH4 measurements at Mondsee.

---

**Table S 1:** biological properties of water samples. Higher values are highlighted column-wise in red, whereas lower values in blue. Abbreviations: pH, power of hydrogen; DO, dissolved oxygen (mg/l); Cond, conductivity (mS m-1); Tur, turbidity (NTU); Temp, temperature (°C); Sal, salinity; Chl.Lab, chloropyll-a content (μg/l), TP, total phosphorus (μg/l); DOC, dissolved organic carbon (μg/l); DN, dissolved nitrogen (μg/l); N_Filter, particulate nitrogen (μg/l); C_Filter, particulate carbon (μg/l); Seston, C:N ratio of suspended particulate matter. Color coding refers to the magnitude for a specific measurement and each parameter (blue low and red high).

| Date | Lake | Abbreviation | pH | DO | Cond | Tur | Temp | Sal | Chl.Lab | TP | DOC | DN | N_Filter | C_Filter | Seston |
| --- | --- | --- | --- | --- | --- | --- | --- | --- | --- | --- | --- | --- | --- | --- | --- |
| 2018-07-11 | Caldonazzo | CAL | 7.74 | 4.15 | 29.1 | 12.2 | 24.2 | 0.1 | 2.45 | 7.79 | 2482.0 | 195.3 | 56.72 | 502.22 | 8.85 |
| 2018-07-12 | Kalterersee | KAL | 7.82 | 3.82 | 46.5 | 16.9 | 25.3 | 0.2 | 3.77 | 9.75 | 3936.0 | 814.6 | 133.82 | 1373.39 | 10.26 |
| 2018-07-17 | Achensee | ACH | 7.92 | 3.89 | 24.9 | 3.2 | 17.0 | 0.1 | 0.66 | 3.20 | 1351.0 | 341.0 | 34.60 | 248.25 | 7.17 |
| 2018-07-18 | Mondsee | MON | 7.91 | 4.24 | 28.7 | 4.1 | 21.4 | 0.1 | 2.48 | 9.10 | 2140.0 | 372.4 | 56.11 | 510.98 | 9.11 |
| 2018-07-19 | Attersee | ATT | 7.90 | 3.90 | 25.0 | 3.8 | 22.6 | 0.1 | 0.61 | 2.10 | 1529.0 | 504.8 | 33.30 | 322.00 | 9.67 |
| 2018-07-24 | Achensee | ACH | 7.93 | 3.84 | 25.3 | 2.8 | 19.4 | 0.1 | 0.64 | 1.80 | 1343.0 | 335.2 | 25.92 | 213.48 | 8.24 |
| 2018-07-25 | Heiterwanger See | HEI | 7.97 | 3.87 | 29.0 | 2.3 | 19.1 | 0.1 | 0.33 | 2.10 | 1048.0 | 324.4 | 18.53 | 99.20 | 5.35 |
| 2018-07-25 | Plansee | PLA | 7.96 | 3.84 | 28.9 | 2.3 | 20.4 | 0.1 | 0.30 | 1.80 | 985.7 | 352.0 | 11.78 | 74.31 | 6.31 |
| 2018-07-30 | Zoggler Stausee | ZOG | 7.95 | 4.39 | 12.1 | 2.3 | 19.8 | 0.0 | 3.18 | 2.92 | 720.4 | 229.6 | 45.30 | 335.39 | 7.40 |
| 2018-08-02 | Achensee | ACH | 7.96 | 4.47 | 25.7 | 3.9 | 21.0 | 0.1 | 0.56 | 2.62 | 1160.0 | 359.3 | 32.19 | 244.37 | 7.59 |
| 2018-08-29 | Lunzer See | LUN | 8.15 | 5.51 | 22.7 | 2.5 | 20.4 | 0.1 | 1.04 | 3.75 | 1628.0 | 581.6 | 27.66 | 221.94 | 8.02 |
| 2018-08-31 | Mondsee | MON | 8.04 | 7.17 | 25.7 | 4.7 | 21.5 | 0.1 | 2.86 | 6.05 | 2204.0 | 227.5 | 48.36 | 525.09 | 10.86 |
| 2018-09-25 | Achensee | ACH | 8.47 | 6.21 | 23.6 | 2.8 | 15.6 | 0.1 | 0.96 | 2.03 | 1503.0 | 324.9 | 25.56 | 203.55 | 7.96 |
| 2018-10-03 | Caldonazzo | CAL | 8.77 | 5.81 | 26.2 | 1.9 | 19.8 | 0.1 | 1.79 | 9.39 | 2497.0 | 201.8 | 42.74 | 249.96 | 5.85 |
| 2018-10-04 | Kalterersee | KAL | 8.64 | 5.96 | 45.0 | 4.1 | 18.7 | 0.2 | 3.53 | 8.15 | 3879.0 | 767.9 | 188.79 | 1770.15 | 9.38 |
| 2018-10-05 | Zoggler Stausee | ZOG | 7.63 | 5.55 | 12.3 | 2.2 | 14.0 | 0.0 | 1.29 | 2.97 | 618.9 | 231.3 | 31.06 | 159.79 | 5.14 |
| 2018-10-16 | Heiterwanger See | HEI | 8.27 | 6.63 | 29.6 | 2.6 | 12.2 | 0.1 | 0.58 | 1.81 | 998.1 | 358.9 | 27.86 | 102.93 | 3.69 |
| 2018-10-16 | Plansee | PLA | 8.44 | 6.46 | 28.6 | 2.3 | 13.8 | 0.1 | 0.61 | 1.03 | 996.0 | 349.4 | 23.78 | 103.49 | 4.35 |
| 2018-10-17 | Kalterersee | KAL | 8.63 | 7.27 | 46.9 | 3.7 | 17.7 | 0.2 | 4.12 | 5.68 | 4197.0 | 807.0 | 120.21 | 1415.88 | 11.78 |
| 2019-05-07 | Kalterersee | KAL | 8.23 | NA | 50.6 | 4.0 | 16.0 | 0.2 | 2.75 | 6.83 | 3413.0 | 1395.0 | 127.63 | 881.70 | 6.91 |
| 2019-06-05 | Mondsee | MON | 8.42 | 5.85 | 29.9 | 3.3 | 20.1 | 0.1 | 0.53 | 4.01 | 1728.0 | 491.9 | 29.49 | 229.82 | 7.79 |
| 2019-06-17 | Kalterersee | KAL | 8.22 | 5.17 | 49.6 | 5.9 | 26.9 | 0.2 | 2.18 | 4.37 | 3298.0 | 1159.0 | 117.64 | 1196.72 | 10.17 |
| 2019-06-19 | Zoggler Stausee | ZOG | 7.63 | 5.61 | 6.0 | 3.4 | 15.2 | 0.0 | 1.40 | 7.39 | 642.2 | 275.2 | 43.05 | 258.18 | 6.00 |
| 2019-06-26 | Mondsee | MON | 8.30 | 4.49 | 29.5 | 4.2 | 26.2 | 0.1 | 0.78 | 4.30 | 1866.0 | 464.2 | 50.12 | 526.86 | 10.51 |
| 2019-07-09 | Zoggler Stausee | ZOG | 7.75 | 5.04 | 9.3 | 3.0 | 16.6 | 0.0 | 0.82 | 3.11 | 531.3 | 266.4 | 40.97 | 205.61 | 5.02 |
| 2019-07-10 | Kalterersee | KAL | 8.36 | 4.94 | 48.0 | 5.6 | 26.9 | 0.2 | 3.04 | 7.64 | 3502.0 | 964.7 | 106.77 | 1020.32 | 9.56 |
| 2019-07-17 | Mondsee | MON | 8.29 | 5.20 | 29.5 | 4.8 | 22.0 | 0.1 | 1.75 | 2.60 | 1938.0 | 447.5 | 37.64 | 524.15 | 13.93 |
| 2019-08-28 | Kalterersee | KAL | 8.39 | 5.43 | 48.4 | 4.1 | 26.2 | 0.2 | 1.79 | 6.87 | 9028.0 | 809.5 | 127.89 | 1270.74 | 9.94 |
| 2019-08-29 | Zoggler Stausee | ZOG | 7.84 | 6.03 | 12.4 | 2.5 | 18.1 | 0.0 | 1.89 | 2.20 | 670.9 | 284.9 | NA | NA | NA |
| 2019-09-04 | Mondsee | MON | 8.30 | 5.00 | 27.5 | 9.4 | 21.4 | 0.1 | 2.91 | 4.16 | 2032.0 | 369.0 | 130.06 | 2274.50 | 17.49 |
| 2019-10-01 | Mondsee | MON | 8.35 | 5.05 | 29.4 | 7.3 | 17.4 | 0.1 | 3.04 | 5.70 | 2105.0 | 394.2 | 94.20 | 1006.56 | 10.69 |
| 2019-10-07 | Kalterersee | KAL | 8.42 | 5.39 | 49.1 | 4.7 | 18.3 | 0.2 | 3.07 | 5.41 | 3740.0 | 912.1 | 57.72 | 1088.30 | 18.86 |
| 2019-10-08 | Zoggler Stausee | ZOG | 7.85 | 4.98 | 14.5 | 2.6 | 12.6 | 0.0 | 1.68 | 2.13 | 456.3 | 252.8 | 13.90 | 245.13 | 17.63 |

**Figure S 1:**
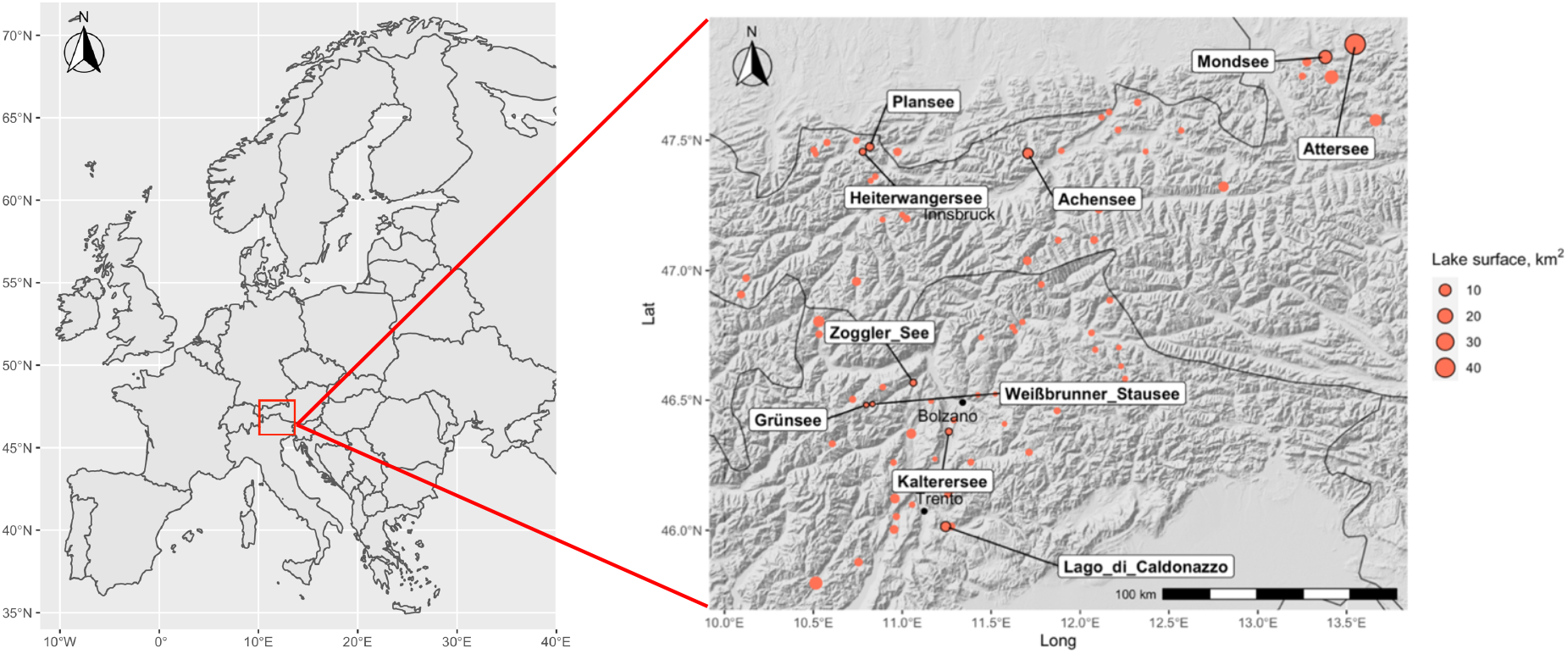
Geographical distribution and area of all lakes in the initial dataset. Those which were sampled during the field campaigns are labeled.

**Figure S 2:**
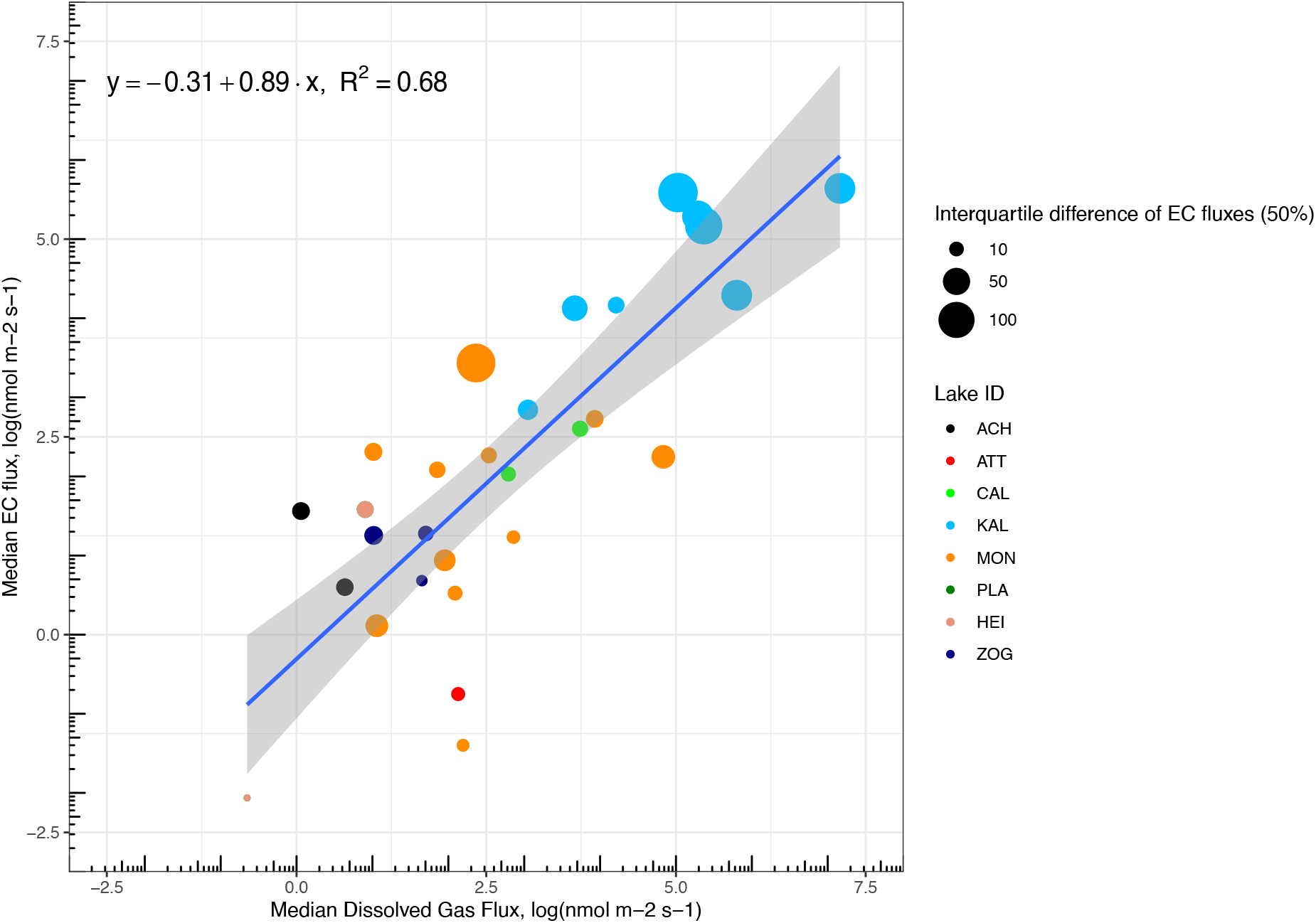
Comparison of EC CH4 fluxes with fluxes estimated from the boundary layer method Spatial variability of EC fluxes is indicated by the size of the symbols (interquartile difference). ACH: Achensee; ATT: Attersee; CAL: Lago di Caldonazzo; HEI: Heiterwangersee; KAL: Kalterersee; MON: Mondsee; PLA: Plansee; ZOG: Zogglersee.

**Figure S 3:**
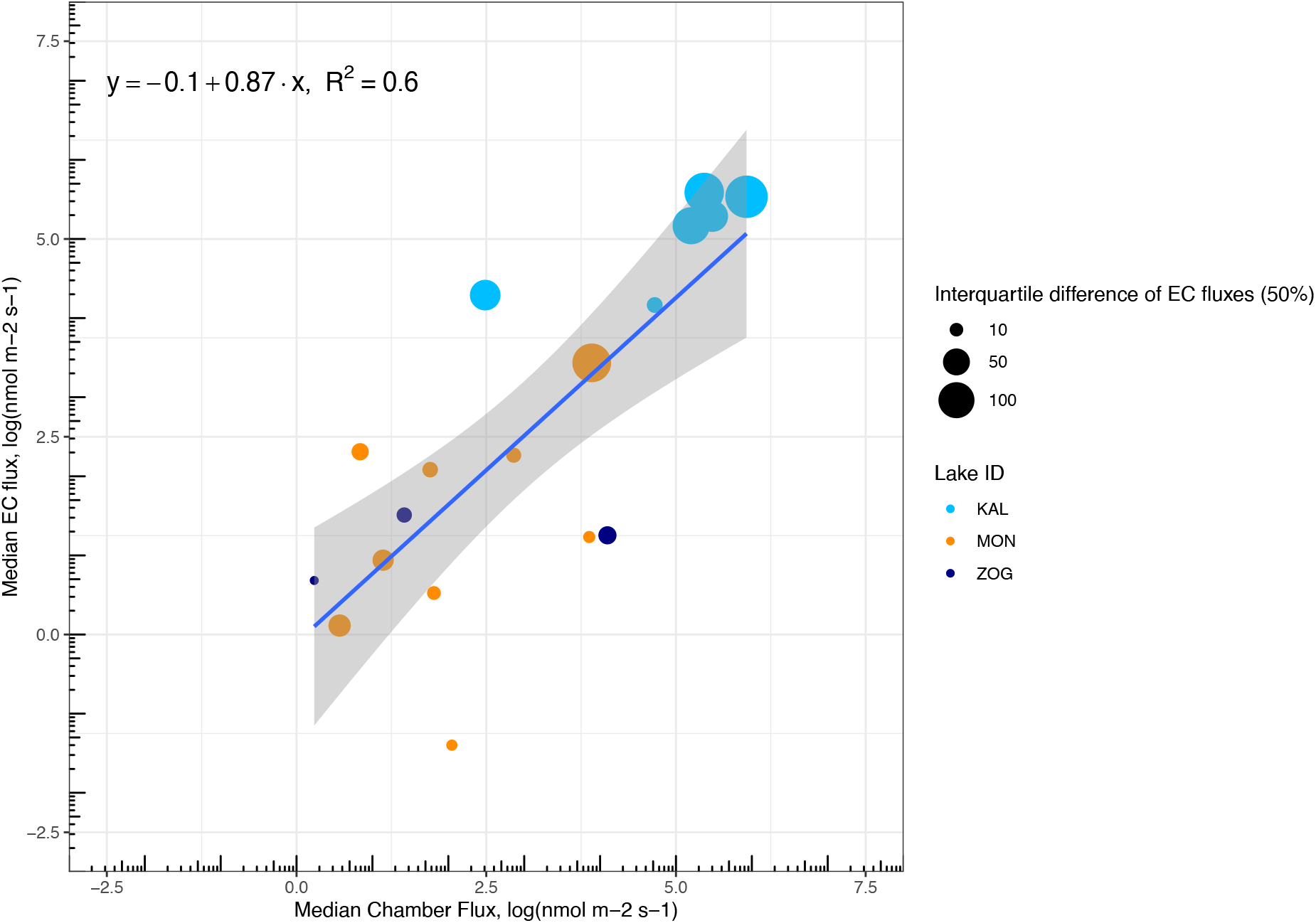
Comparison of EC CH4 fluxes with fluxes estimated with chambers. Spatial variability of EC fluxes is indicated by the size of the symbols (interquartile difference). KAL: Kalterersee; MON: Mondsee; ZOG: Zogglersee.

